# A widespread thermodynamic effect, but maintenance of biological rates through space across life’s major domains

**DOI:** 10.1101/274944

**Authors:** Jesper G. Sørensen, Craig R. White, Grant A. Duffy, Steven L. Chown

## Abstract

For over a century, temperature compensation (maintenance of biological rates with changing temperatures) has remained controversial. An alternative idea, that fitness is greater at higher temperatures (the thermodynamic effect), has gained increasing traction, and is being used to understand large-scale biodiversity responses to environmental change. Yet evidence in favour of each of these contrasting hypotheses continues to emerge. In consequence, the fundamental nature of organismal thermal responses and its implications remain unresolved. Here we investigate these ideas explicitly using a global dataset of 619 observations of four categories of organismal performance, spanning 14 phyla and 403 species. In agreement with both hypotheses, we show a positive relationship between the temperature of maximal performance rate (T_opt_) and environmental temperature (T_env_) for all traits. Next we demonstrate that relationships between T_env_ and the temperature of maximal performance rate (U_max_) are rarely significant and positive, as expected if a thermodynamic effect predominates. By contrast, a positive relationship between T_opt_ and U_max_ is always present, but markedly weaker than theoretically predicted. These outcomes demonstrate that while some form of thermodynamic effect exists, ample scope is present for biochemical and physiological adaptation to thermal environments in the form of temperature compensation.

## Introduction

All organisms are exposed to variation in ambient temperature. Such variation typically has direct effects on the physiology and population dynamics of ectotherms, ultimately exerting a marked influence on range size and dynamics (1-3). In consequence, ectothermic animals and plants exhibit a wide range of responses to modulate the effects of ambient temperature variation (4-6). Among their adaptive responses, temperature compensation has proven especially controversial. Also known as metabolic cold adaptation (7), the Krogh effect (8), or metabolic compensation (9), temperature compensation refers to the maintenance of biological rates in the face of a temperature change (10-12). Initially proposed on the basis of empirical evidence and the theoretical notion that rate maintenance, especially under low temperature conditions, would result in maintenance of fitness (13-15), the idea has become controversial on both theoretical and empirical grounds. The controversy has been most prominent for metabolic rate conservation, with the theoretical counterargument being that because metabolic rate represents a cost (of maintenance) to an organism, conservation thereof, in the face of an opportunity for reduction, should not be beneficial (11). Empirical evidence, typically from measurements of standard or resting metabolic rates across a range of biological levels, has come out both in favour of and against temperature compensation (3, 5, 9, 15-22).

One line of evidence that has been especially effective in questioning the temperature compensation hypothesis is the discovery and description of a thermodynamic effect (23). Sometimes also known as the ‘warmer is better’ hypothesis, the idea encompasses both sound theoretical reasons and evidence for a relationship between the optimum temperature of a process and the maximal rate of that process (Figure 1). In other words, because rates proceed faster at higher ambient, and therefore by association for many ectotherms, higher organismal temperatures, fitness should always be higher at higher temperatures, acknowledging that upper thermal limits to performance exist for all organisms (24-25). The strongest evidence for the thermodynamic effect comes from population growth rates in insects, with suggestions that it applies to performance traits in ectotherms generally (26-28). Across the 65 insect species examined by Frazier et al. (26), the thermodynamic effect was found to be even stronger than predicted by theory (29), suggesting that relatively warm environments have the highest fitness benefits for organisms. In turn, these findings have also been used to explain the slow life histories of polar organisms (21).

**Figure 1.**
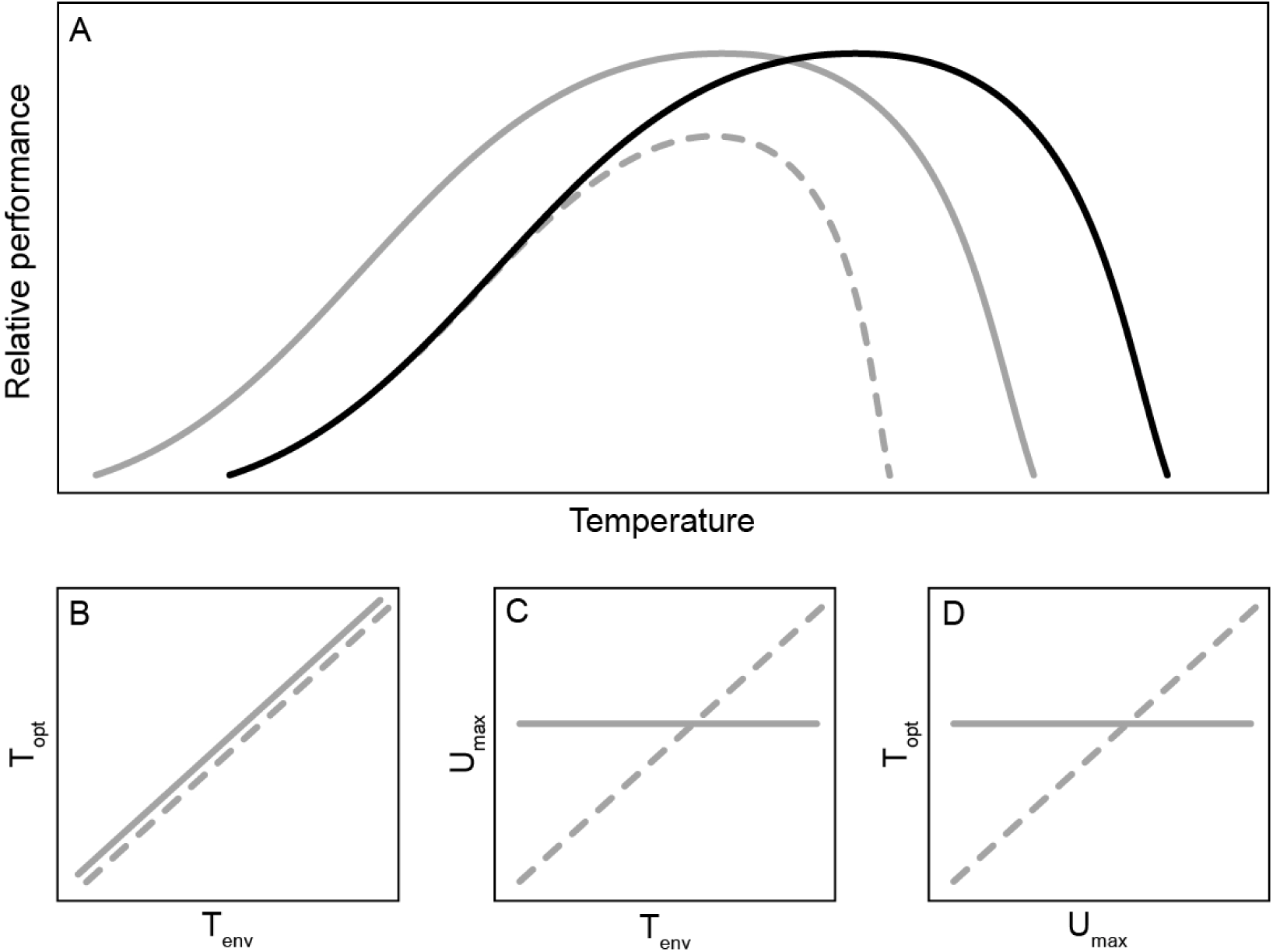
Conceptual figure showing expected relationships under either the temperature compensation or thermodynamic effect hypothesis. The relative performance of a given trait (a) is expected to increase with temperature until peak performance (U_max_) is achieved at the optimum temperature (T_opt_), after which performance declines (solid black line). In colder climates, the temperature compensation hypothesis predicts that the relationship between relative performance and temperature will shift such that U_max_ occurs at a lower T_opt_, but remains equal to that observed in warmer climates if fullcompensation is achieved (solid grey line). Alternatively, the thermodynamic effect hypothesis predictsthat in colder climates U_max_ will not only occur at a lower T_opt_, but will also be lower than that observed inwarmer climates (dashed grey line). Panels below show the expected relationships between (b)environmental temperature (T_env)_ and T_opt_, (c) T_env_ and U_max_, (d) U_max_ and T_opt_, respectively under thetemperature compensation (solid lines) and thermodynamic effect (dashed lines) hypotheses. Bothhypotheses predict a positive correlation between T_env_ and T_opt_ (b). However, the temperaturecompensation hypothesis predicts that T_opt_ will be independent of T_env_ while a positive relationship isexpected under the thermodynamic effect hypothesis (c). Likewise, U_max_ is expected to be independent of T_opt_ under temperature compensation, while the thermodynamic effect hypothesis predicts a positiverelationship (d).

Despite this evidence for a thermodynamic effect, several studies continue to find empirical support for temperature compensation. For example, in plants, much evidence has been found for maintenance of respiration rate across a broad range of temperatures (5-6, 9). In other groups, contrasting empirical outcomes continue to be published (22, 30), with little indication of a developing consensus (8, 21). In consequence, despite the existence of the idea of temperature compensation for a century (13), and strong theoretical and empirical bases for the thermodynamic effect (29, 31), how these contrasting ideas are related, and which might provide the strongest explanation for the evolution of biological rates in response to temperature variation across the globe remains at best unclear. Moreover, explanations also seem to differ in their support across different groups of organisms and from different environments (11), and often with little comparison among taxa (compare e.g. 17, 21, 32), though with notable exceptions (9, 3). Yet at the same time, the expectations from these competing hypotheses are variously being used as the basis to understand diversity variation globally (34) and the extent to which changes in this diversity might occur as a consequence of anthropogenic warming (35-36).

Here we seek to resolve these long-standing and important (11) contrasting ideas by examining optimum temperature and rates at those optima for a suite of biological functions across much of life’s ectotherm diversity and at a global scale. Rather than treating major taxa and organisms from terrestrial and aquatic habitats separately, we use phylogenetic mixed models to investigate the extent to which both habitat and phylogenetic signal influence the relationships between optimum temperature and rates of biological functions at that temperature, and subsequently the ways in which both optimum temperature and maximum rates vary with temperature across the planet. We focus on rates of development, growth, locomotion and photosynthesis, which are expected to be correlated with fitness (26), but we avoid investigation of metabolic rates (or respiration rate for plants, e.g. 9). We do so because few animal ectotherm metabolic rate investigations provide measured values for maximal rates and the temperatures thereof (U_max_ and T_opt_ in the terminology of 37).

Our analysis uses information from 619 observations, spanning 14 phyla, 75 orders, 300 genera and 403 species. By contrast with previous comprehensive analyses of the slope of the relationship between rate and temperature (e.g. 33), we are concerned here with optimum rates (U_max_) and the temperatures at which they occur (T_opt_). We test explicitly three predictions of the temperature compensation and thermodynamic effect hypotheses. First, if either of these hypotheses holds, a positive relationship between T_opt_ and a measure of environmental temperature (T_env_) during the maximal activity period of the organism should be found (Figure 1), assuming that some form of thermal adaptation (or coadaptation) is typical of ectotherms (38-40). Absence of a relationship might indicate some form of performance constraint (41). Second, the relationship between U_max_ and T_env_ should be positive in the case of the predominance of a thermodynamic effect, but absent or weak in the case of temperature compensation (40, 42). Third, U_max_ and T_opt_ should be positively related in the case of a pronounced thermodynamic effect, but weak or absent where temperature compensation predominates (26). More specifically, when U_max_ is plotted against the inverse of optimum body temperature, the thermodynamic effect hypothesis suggests that the slope of the line should provide an estimate of activation energy of 0.6 to 0.7 eV or perhaps steeper (23, 26, 29).

## Methods

We compiled data from published literature on optimal temperature and maximal performance (T_opt_ and U_max_ (*sensu* 37) for whole organismal traits expected to be closely related to fitness including rates of photosynthesis, growth, development and locomotion performance. Many published studies are available for these traits, making it possible for the database to cover the majority of the world and a diverse range of taxonomic groups and habitats to gain general insight. In additional to original papers, recent compilations of data and their reference lists were also searched (26, 28, 32, 43-46). The search ended on January 1^st^ 2016. We only accepted records where measured estimates of performance were undertaken beyond the measured maximal performance (i.e. T_opt_ and U_max_). Performance curves where maximal performance was only estimated by models were not included. For development rates, however, high temperatures leading to no development were accepted as a data point above maximal performance. We included the full taxonomy of all organisms as given by the primary publication, and adjusted for synonymy where appropriate based on online repositories (such as www.algaebase.org or www.gbif.org). The analyses were done according to the species lists as generated by the online tree of life (47). The geographical origin of the investigated population of each species (and for each trait where the locations differed among traits) was taken from the primary literature whenever possible. When the origin of an investigated population was not available from primary literature, the origin was estimated using data from the Global Biodiversity Information Facility (GBIF). Median latitude and longitude was extracted from GBIF occurrence records using the ‘rgbif’ (48) and ‘spocc’ (49) packages in R (50) and used for that species. In cases where GBIF records were lacking the origin was estimated from other sources (described for each record in the database, Table S17). For locomotion we included ln-transformed body length as a covariate, and for developmental rates we included ln-transformed dry mass as a covariate because of significant allometry of these traits (see results). Snout-vent length (for reptiles and anurans) and body length (for fish and invertebrates) were obtained from the original literature or estimated from other sources when not available (described for each record in the database, Table S17). Dry mass estimates were sourced from the original literature when given or inferred from length or fresh mass measured available using specific relationships given by Hodar (51) and Ganihar (52). In all cases the sources and relationships used to generate dry mass estimates are given in the database.

Data were analysed using phylogenetic mixed models (53-55), which were implemented in the ‘ASReml-R’ v3.0 (56) package of R v3.0.2 (57), with inverse relatedness matrices calculated from phylogenetic covariance matrices using the ‘MCMCglmm’ package v2.21 (58). The phylogeny used for analysis was drawn from a comprehensive tree of life, accessed using the ‘rotl’ v0.5 package of R v3.2.2 (47, 59). In addition to the 619 observations that were analysed, a further 319 records for 80 species were excluded from the analysis; some of these could not be matched to the online tree of life, and so were not considered further. Six extremely high maximum rates for growth of Actinobacteria from the Luna-2 cluster (60) and one extremely high growth rate for *Chlorella pyrenoidosa* (61) exerted high leverage on the data and were excluded on these grounds. Twenty-two records were removed because they could not be matched to climate data, one record was removed because the temperature of the warmest quarter was less than 0 °C; the remaining records were excluded because they data were presented in units that could not be reasonably converted to match the majority of the remaining data.

Environmental temperature (T_env_) at the site of geographical origin (see above) for each record was calculated as the mean temperature of the warmest quarter using monthly (January 2001 – December 2016) daytime data from the MODIS Land Surface Temperature dataset (MOD11C3 v6;doi:10.5067/MODIS/MOD11C3.006; 0.05° spatial resolution). Seasonality at each site was calculated as the difference between the mean temperature of the warmest quarter and the mean temperature of the coldest quarter, also calculated from the MODIS Land Surface Temperature dataset. These data were downloaded and analysed using the ‘MODIS’ (62), ‘raster’ (63), and ‘xts’ (64) packages in R (50).

Phylogenetic mixed models were selected over the more commonly used methods of independent contrasts (65) and phylogenetic generalised least squares (66) because the former can formally incorporate non-independence associated with phylogenetic relatedness as well as non-independence associated with multiple measurements of single species. Multiple measurements were relatively uncommon in the data sets for locomotion, growth, and development, where 73%, 88%, and 91% of species were represented by only one measurement, respectively, though a small number of species were represented by many measurements (up to ten measurements per species for locomotion, up to 14 measurements per species for growth, and up to five measurements per species for development).Multiple measurements are more common in the data for photosynthesis, where 33% of species have one measurement, 49% of species have two measurements and the remainder have three-to-eight measurements. Phylogenetic mixed models are an analogue of the animal model from quantitative genetics, which partitions phenotypes of related individuals into heritable (additive genetic) and non-heritable components to estimate inter-specific variances and covariances between traits (55). The significance of fixed effects was tested using Wald-type *F*-tests with conditional sums of squares and denominator degrees of freedom calculated according to Kenward and Roger (67). Phylogenetic heritability, a measure of phylogenetic non-independence equivalent to Pagel’s (68) *λ* (55), was estimated as the proportion of variance attributable to the random effect of phylogeny. Approximate standard errors for the estimate of phylogenetic heritability was calculated using the R ‘pin’ function (69).

## Results

We used phylogenetic mixed models to investigate the relationship between optimum temperature (T_opt_) and environmental temperature (T_env_), measured here as mean temperature of the warmest quarter (derived from the Moderate Resolution Imaging Spectroradiometer, MODIS, https://modis.gsfc.nasa.gov/) of the collection locality of the species concerned (see Methods). The results demonstrated a positive relationship, though with much variation, for development rate, and no relationship between T_opt_ and T_env_ for growth rate, locomotion rate, and photosynthetic rate (Figure 2). Interaction terms in these models were always non-significant. Thus, only models with additive combinations of main effects are presented. For all traits a strong phylogenetic signal was detected (Phylogenetic heritability □ 0.82 – 0.98; Tables S1-S4).

**Figure 2.**
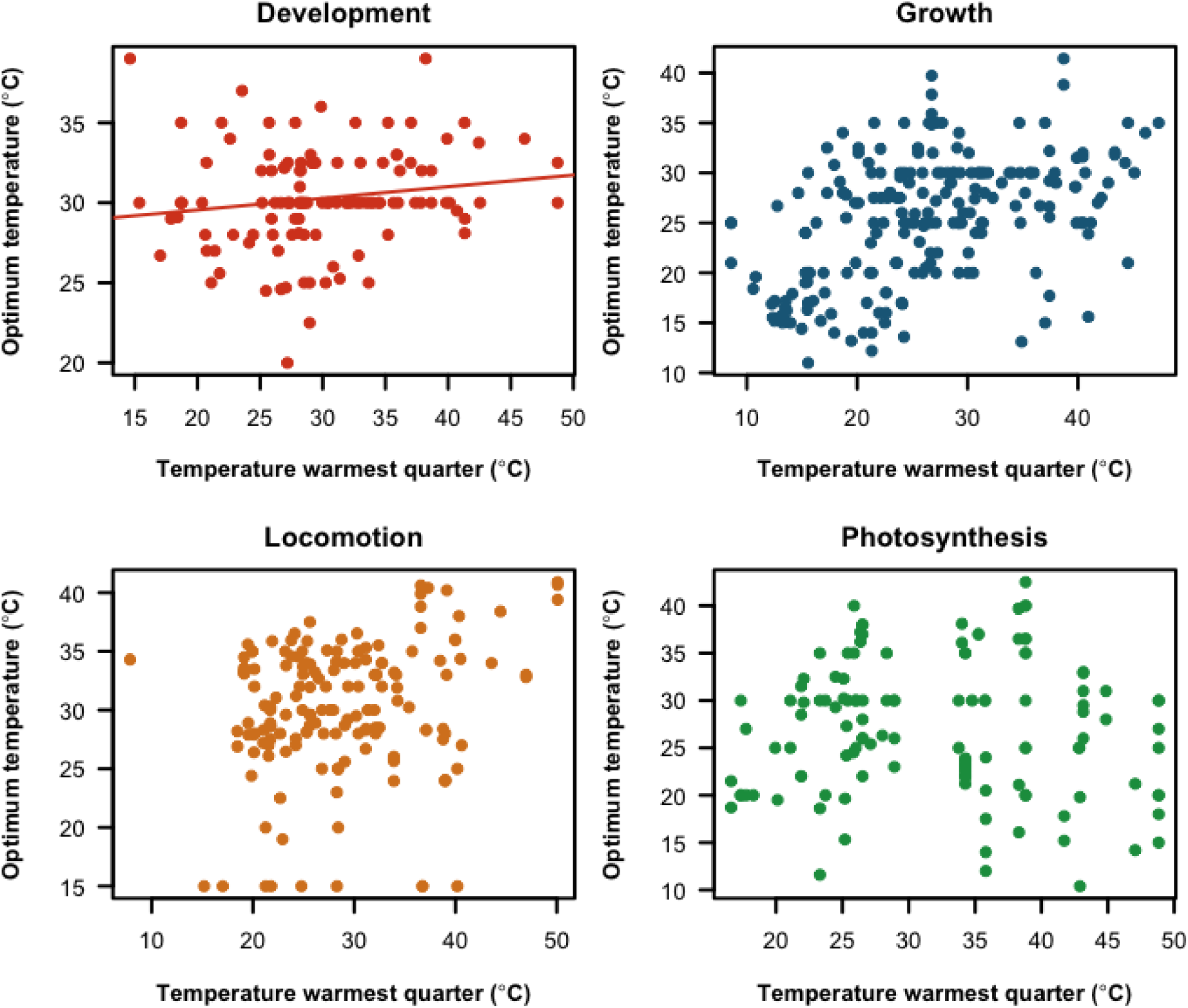
Relationship between mean temperature of the warmest quarter of the year (as a measure of Tenv, °C) and the optimum temperature (Topt, °C) for rates of development (d^−1^), growth (% d^−1^), locomotion (cm s^−1^) and photosynthesis (μmol m^−2^ s^−1^), respectively. Statistical outcomes are provided in Tables S1-S4. Solid lines depict significant relationships from phylogenetic mixed models testing for effects of T_env_ on T_opt_ (Table S1).

In the case of the relationship between natural log-transformed maximal performance (U_max_) and our measure of T_env_, no relationship was found for any of the performance traits (Figure 3). Again, interaction terms were never significant and the phylogenetic signal was strong (Phylogenetic heritability □⁏0.76 – 0.98; Tables S5-S8).

**Figure 3.**
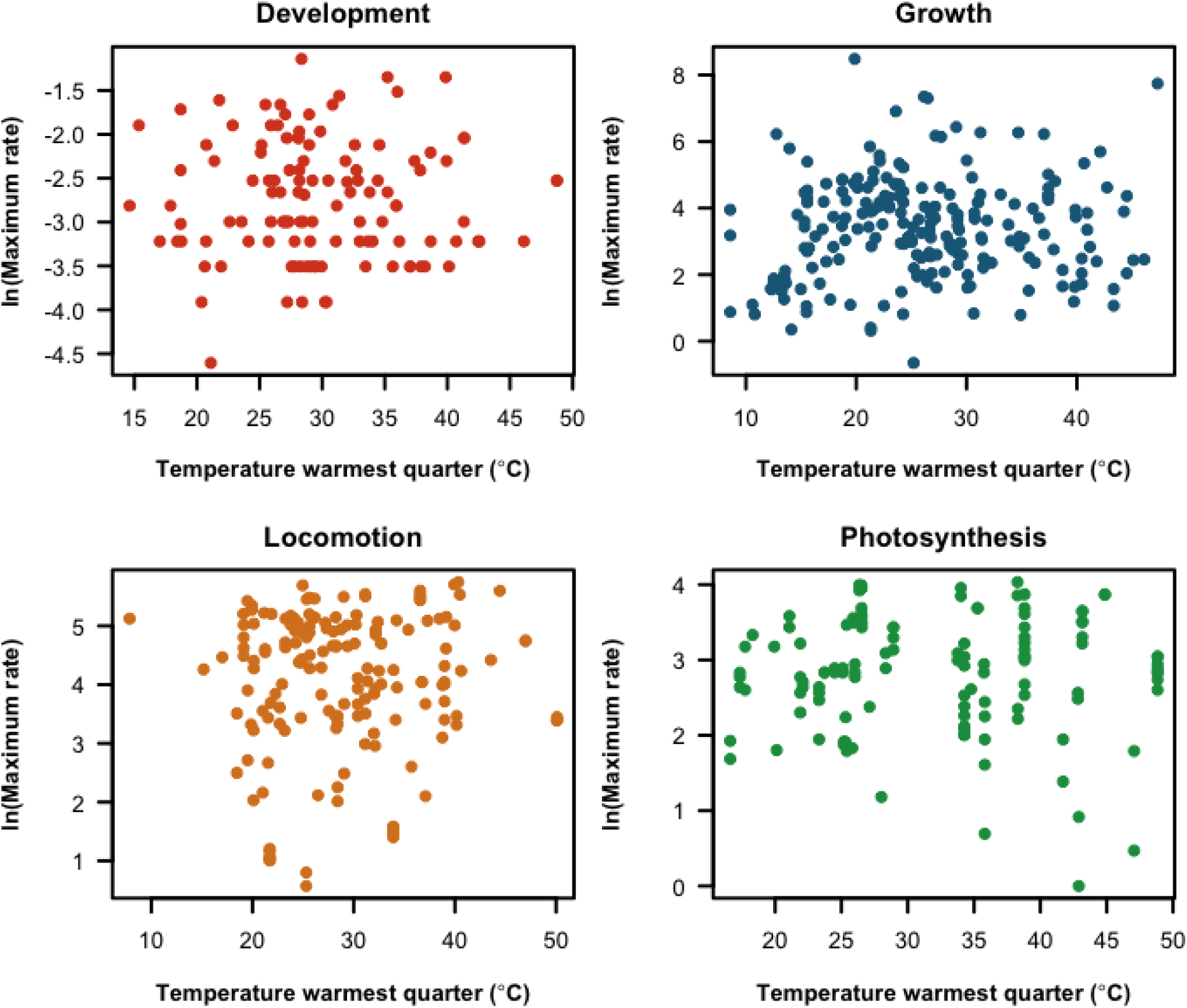
Relationship between mean temperature of the warmest quarter of the year (as a measure of Tenv, °C) and the natural log-transformed maximum rate (Umax) for rates of development (d^−1^), growth (% d^−1^), locomotion (cm s^−1^), and photosynthesis (μ**mol m**^**-2**^ **s**^**-1**^), respectively. Statistical outcomes are provided in Tables S5-S8.

By contrast with these variable outcomes, a positive relationship between maximal performance (U_max_) and optimal temperature (T_opt_) was characteristic of all the traits examined in models that considered only main effects without interaction terms: development rate, growth rate, locomotion speed and photosynthetic rate (Figure 4, Tables S9-S12), again with much variation about the central tendencies. In the full factorial models, however, phylum and T_opt_ showed significant interactions (Table S13) for growth rate, as did T_opt_ and phylum for locomotion rate (Table S14). Data for growth rate were therefore further subdivided by phylum (Figure S1), but there were locomotion data for too few species of arthropod to formally estimate model parameters for this phylum alone. Significant positive relationships between U_max_ and T_opt_ characterised the subdivided datasets (Table S15). When converted to activation energy, values ranged between 0.16 and 0.68 eV, with a mean of 0.37 ± 0.08 [SE] eV, which is significantly different from the value of 0.60 eV predicted from theory (26) (*t*_5_ = −2.93, p = 0.03), but not from 0.54 eV (*t*_5_ = −2.16, p = 0.08), previously a minimum empirical value (23).

**Figure 4.**
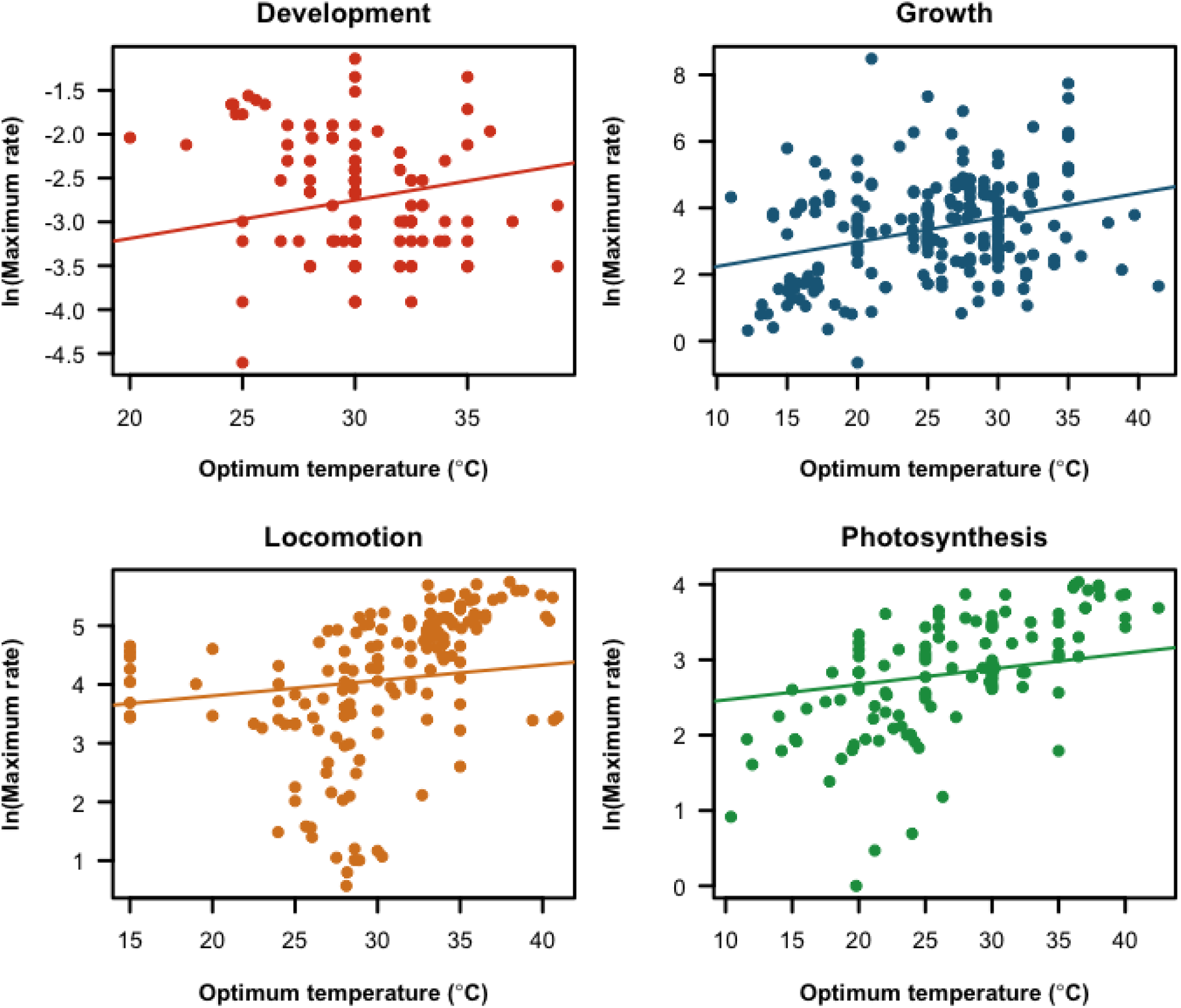
Relationship between optimum temperature (Topt, °C) and the natural log-transformed maximum rate (Umax) for rates of development (d^−1^), growth (% d^−1^), locomotion (cm s^−1^), and photosynthesis (μmol m^−2^ s^−1^), respectively. Solid lines depict significant relationships from phylogenetic mixed models testing for significant effects of T_opt_ on U_max_ (Tables S9-S12).

## Discussion

Understanding the nature of and potential limitations characterising physiological and biochemical adaptation to temperature is a fundamental question in organismal biology (4, 12, 70). Moreover, what form such adaptation might take, as reflected in the relationship between temperature and biological rates, has important implications for interpreting the responses of organisms to changing environments, including the influences of global climate change (6, 20, 35-36, 40). For example, if the thermodynamic effect predominates in the relationship between U_max_ and T_opt_, rising temperatures might prove largely beneficial for ectotherms except perhaps in the tropics (though see 44) because biochemical constraints are reduced. By contrast, if some form of compensation is more common, changing temperature regimes may have less of an effect on performance (9, 11, 36). Thus, which of these major relationships between U_max_ and T_opt_ predominate is of both fundamental and applied significance.

Previous examinations of the relationship between U_max_ and T_opt_ have come out strongly in favour of the thermodynamic effect hypothesis (23, 26-28), with activation energies either being within the predicted range of 0.6 to 0.7 eV (23), or larger, implying a stronger thermodynamic effect than theoretically predicted (26). Based on a much larger suite of data, spanning a wide range of localities, habitats and taxa, and several key performance traits, we also find that the thermodynamic effect is generally supported for the relationship between U_max_ and T_opt_. In contrast with previous investigations, however, we find this effect (on average an activation energy of 0.37 ± 0.08 eV) much weaker than proposed by theory or previously found empirically (i.e. 0.6-0.7 eV, or 0.54 to 0.97 eV) (23, 26). Thus, while a thermodynamic effect is general, it is not pronounced.

The difference between this finding and that of previous studies might owe in part to the inclusion of a specific plant performance trait, photosynthetic rate, in the current investigation. The activation energy value for this trait was lowest of all of the significant values (0.16 eV); with the value for photosynthetic rate excluded, the mean activation energy increases to 0.41 ± 0.08 and is not significantly different from 0.54 (*t*_4_ = −1.57, p = 0.19) or 0.60 (*t*_4_ = −2.30, p = 0.08). This change does, however, point to a further explanation for the different outcomes between our study and others. The consideration of organisms from a wide variety of environments, which represent several life history types and trophic groups is likely to mean much larger variation in the way U_max_ and T_opt_ are related, and how these traits are related to environmental temperature (11, 33-34, 40). For example, owing to their restricted movement capability, plants may be expected to show a much greater level of thermal compensation than ectotherm animals, which can behaviourally select among a diversity of thermal microenvironments available to them in any given larger setting (4, 71). Indeed, temperature compensation of respiration rates in plants of several different groups is commonly found (5, 9, 20). The same preponderance of compensation might be expected in aquatic versus non-aquatic groups, given the thermal inertia of aquatic environments (72). In the one case where we were able to draw such an explicit contrast – for locomotion speed in aquatic versus non-aquatic chordates (Table 1; Figure S1) – the variation is in the direction predicted, with no significant relationship between U_max_ and T_opt_ for the aquatic group. Nonetheless, for metabolic rate variation the reverse seems to be true, with compensation being less commonly found in aquatic marine groups than in terrestrial species (11-12).

The relatively weak relationship between U_max_ and T_opt_ does point to the fact that some form of thermal compensation is characteristic of all the organisms we examined, in keeping with long-standing contentions about the importance thereof (10, 13). The typical absence of a relationship between U_max_ and T_env_ here also supports this contention, because the absence of a relationship is predicted by the hypothesis of temperature compensation (42). For photosynthesis rate, the outcome is clearly in keeping with findings for plants, and in particular for respiration rate, where compensation is well documented (5-6, 9, 20). For the other traits, and especially in animal ectotherms, the findings contrast with those from the broader thermal performance literature (4, 28). The variability around the T_env_ and T_opt_ relationship in the traits excluding photosynthesis is also perhaps surprising, although here positive relationships for development rate and for locomotion speed are in keeping with previous work (26). Nonetheless, our results demonstrate that temperature compensation is more commonplace than previously estimated for animals.

Several caveats should be borne in mind, however. First, a mismatch between T_env_ and the peak characteristics of the performance curve – T_opt_ and U_max_ – might be expected because such differences,especially between T_env_ and T_opt_ could be an adaptive response to environmental seasonality (40). In this case, the difference between T_env_ and T_opt_ should be strongly related to a measure of environmental seasonality, with a potential difference between tropical and non-tropical organisms. We tested for such an effect of seasonality and found that the strength of the effect varied among traits and phyla, with significant relationships between seasonality and the difference between T_env_ and T_opt_ found only for locomotion and photosynthesis rates (Table S16, Figure S2). The latter accords well with recent finding that terrestrial net primary production is better predicted by growing season length than by latitude or temperature (73). Thus, some adaptive response to seasonality may be occurring in these traits, and deserves further consideration. Second, we calculated T_env_ as mean temperature of the warmest quarter from the collection locality of the population investigated (see Methods). This may not fully represent the thermal environment typical of the organisms collected, though it is likely a better estimate of temperature when most organisms are actively growing and developing than mean annual temperature (74). Moreover, the temperature estimate used can have an effect on the form of the relationship between a trait and temperature (75). The estimates of relationships between T_env_ and performance-related traits provided here differ, however, from those made for activation energy of traits in other studies (e.g. 33-34). In those studies, the temperature dependence of the traits is estimated not from comparisons of T_opt_ or U_max_ across species from different environments, but rather from trait values at a given range of experimental temperatures leading up to and moving away from T_opt_ within a given species.

Given these outcomes, it is clear that while some form of thermodynamic effect exists, ample scope is present for biochemical and physiological adaptation in the form of temperature compensation. Indeed, the overriding influence seems to be one of biochemical and physiological adaptation, at least for the traits examined here, so vindicating earlier views on the significance of such adaptation (10, 13, 70, 76). Much variation exists, however, within and among traits, and among taxa and environments. Such variation would have to be considered when using these general relationships to forecast the broader implications of environmental change, as has become clear from related studies of the thermal dependence of performance-related traits (33-34). To some extent the variation seen may also explain the many contrary findings in the literature to date. In the case of assessments based on metabolic rate of animal ectotherms, which have often dominated the animal literature, much of the debate on the existence of compensation (5, 11, 15-19, 21-22) might, however, be overcome by trait assessments which include the full performance curve providing empirical estimates of T_opt_ and U_max_, as is done for plants (e.g. 9),rather than just on the increasing side of the curve.

## Competing interests

The authors declare no competing interests.

## Authors’ contributions

JGS and SLC designed the study and collected the data from the literature. GAD provided input to the design of the study, and prepared environmental data and the conceptual figure. CRW performed the analyses and associated figures. JGS and SLC prepared the first draft of the ms, and all authors contributed to the final version.

## ACKNOWLEDGMENTS

We thank Johannes Overgaard and Lesley Alton for comments on a previous version of the manuscript. Research funding for this project was provided by a Sapere Aude DFF-Starting grant from The Danish Council for Independent Research | Natural Sciences and a sabbatical grant from the Aarhus University Research Foundation (AUFF) to JGS, and by the Australian Research Council through ARC DP170101046 to SLC and FT130101493 to CRW.

## References

1. Cossins AR, Bowler K. 1987 Temperature Biology of Animals. Chapman and Hall, New York.

2. Gaston KJ. 2009 Geographic range limits: achieving synthesis. Proc R Soc B 276, 1395–1406.

3. Overgaard J, Kearney MR, Hoffmann AA. 2014 Sensitivity to thermal extremes in Australian Drosophila implies similar impacts of climate change on the distribution of widespread and tropical species. Global Change Biol 20, 1738–1750.

4. Angilletta MJ. 2009 Thermal Adaptation. A Theoretical and Empirical Synthesis. Oxford University Press, Oxford.

5. Atkin OK, Bloomfield KJ, Reich PB, Tjoelker MG, Asner GP, Bonal D, Bönisch G, Bradford MG, Cernusak LA, Cosio EG et al. 2015 Global variability in leaf respiration in relation to climate, plant functional types and leaf traits. New Phytol 206, 614–636.

6. Heskel MA, O’Sullivan OS, Reich PB, Tjoelker MG, Weerasinghe LK, Penillard A, Egerton JJG, Creek D, Bloomfield KJ, Xiang J et al. 2016 Convergence in the temperature response of leaf respiration across biomes and plant functional types. Proc Natl Acad Sci USA 113, 3832–3837.

7. Clarke A. 1980 A reappraisal of the concept of metabolic cold adaptation in polar marine invertebrates. Biol J Linn Soc 14, 77–92.

8. Gaston KJ, Chown SL, Calosi P, Bernardo J, Bilton DT, Clarke A, Clusella-Trullas S, Ghalambor CK, Konarzewski M, Peck LS et al. 2009 Macrophysiology: A Conceptual Reunification. Am Nat 174, 595–612.

9. Padfield D, Lowe C, Buckling A, Ffrench-Constant R, Student Research Team, Jennings S, Shelley F, Ólafsson JS, Yvon-Durocher, G. 2017 Metabolic compensation constrains the temperature dependence of gross primary production. Ecol Lett 20, 1250–1260.

10. Hazel JR, Prosser CL. 1974 Molecular mechanisms of temperature compensation in poikilotherms. Physiol Rev 54, 620–677.

11. Clarke A. 2003 Costs and consequences of evolutionary temperature adaptation. Trends Ecol Evol 18, 573–581.

12. Clarke A. 2017 Principles of Thermal Ecology. Temperature, Energy and Life. Oxford University Press, Oxford.

13. Krogh A. 1916 The Respiratory Exchange of Animals and Man. Longman, London.

14. Sømme L, Block W. 1991 Adaptations to alpine and polar environments. Insects at Low Temperature, eds Lee RE, Denlinger DL. Chapman & Hall, London, pp 318–359.

15. Chown SL, Gaston KJ. 1999 Exploring links between physiology and ecology at macro-scales: the role of respiratory metabolism in insects. Biol Rev 74, 87–120.

16. Clarke A, Johnston NM. 1999 Scaling of metabolic rate with body mass and temperature in teleost fish. J Anim Ecol 68, 893–905.

17. Addo-Bediako A, Chown SL, Gaston KJ. 2002 Metabolic cold adaptation in insects: a large-scale perspective. Funct Ecol 16, 332–338.

18. Steffensen JF. 2002 Metabolic cold adaptation of polar fish based on measurements of aerobic oxygen consumption: fact or artefact? Artefact! Comp Biochem Physiol A 132, 789–795.

19. White CR, Alton LA, Frappell PB. 2012 Metabolic cold adaptation in fishes occurs at the level of whole animal, mitochondria and enzyme. Proc R Soc B 279, 1740–1747.

20. Padfield D, Yvon-Durocher G, Buckling A, Jennings S, Yvon-Durocher G. 2016 Rapid evolution of metabolic traits explains thermal adaptation in phytoplankton. Ecol Lett 19, 133–142.

21. Peck LS. 2016 A cold limit to adaptation in the sea. Trends Ecol Evol 31, 13–26.

22. Alton LA, Condon C, White CR, Angilletta MJ, Jr. 2017 Colder environments did not select for a faster metabolism during experimental evolution of Drosophila melanogaster. Evolution 71, 145–152.

23. Savage VM, Gillooly JF, Brown JH, West GB, Charnov EL. 2004 Effects of body size and temperature on population growth. Am Nat 163, 429–441.

24. Pörtner HO. 2002 Climate variations and the physiological basis of temperature dependent biogeography: systemic to molecular hierarchy of thermal tolerance in animals. Comp Biochem Physiol A 132, 739–761.

25. Araújo MB, Ferri-Yáñez F, Bozinovic F, Marquet PA, Valladares F, Chown SL. 2013 Heat freezes niche evolution. Ecol Lett 16, 1206–1219.

26. Frazier MR, Huey RB, Berrigan D. 2006 Thermodynamics constrains the evolution of insect population growth rates: “Warmer is better”. Am Nat 168, 512–520.

27. Knies JL, Kingsolver JG, Burch CL. 2009 Hotter is better and broader: Thermal sensitivity of fitness in a population of bacteriophages. Am Nat 173, 419–430.

28. Angilletta MJ, Huey RB, Frazier MR. 2010 Thermodynamic effects on organismal performance: Is hotter better? Physiol Biochem Zool 83, 197–206.

29. Gillooly JF, Brown JH, West GB, Savage VM, Charnov EL. 2001 Effects of size and temperature on metabolic rate. Science 293, 2248–2251.

30. Zhu W, Zhang H, Li X, Meng Q, Shu R, Wang M, Zhou G, Wang H, Miao L, Zhang J et al. 2016 Cold adaptation mechanisms in the ghost moth Hepialus xiaojinensis: Metabolic regulation and thermal compensation. J. Insect Physiol. 85, 76–85.

31. Clarke A. 2004 Is there a universal temperature dependence of metabolism? Funct Ecol 18, 252–256.

32. Cavicchioli R. 2016 On the concept of a psychrophile. ISME J 10, 793–795.

33. Dell AI, Pawar S, Savage VM. 2011 Systematic variation in the temperature dependence of physiological and ecological traits. Proc Natnl Acad Sci USA 108, 10591–10596.

34. Dell AI, Pawar S, Savage V. 2014 Temperature dependence of trophic interactions are driven by asymmetry of species responses and foraging strategy. J Anim Ecol 83, 70–84.

35. Dillon ME, Wang G, Huey RB. 2010 Global metabolic impacts of recent climate warming. Nature 467, 704–706.

36. Seebacher F, White CR, Franklin CE. 2015 Physiological plasticity increases resilience of ectothermic animals to climate change. Nature Climate Change 5, 61–66.

37. Gilchrist GW. 1995 Specialists and generalists in changing environments. I. Fitness landscapes of thermal sensitivity. Am Nat 146, 252–270.

38. Huey RB, Bennett AF. 1987 Phylogenetic studies of coadaptation: preferred temperatures versus optimal performance temperatures of lizards. Evolution 41, 1098–1115.

39. Angilletta MJ, Niewiarowski PH, Navas CA. 2002 The evolution of thermal physiology in ectotherms. J Therm Biol 27, 249–268.

40. Amarasekare P, Johnson C. 2017 Evolution of thermal reaction norms in seasonally varying environments. Am Nat 189, E31–E45.

41. Makarieva AM, Gorshkov VG, Li B-L, Chown SL, Reich PB, Gavrilov VM. 2008 Mean mass-specific metabolic rates are strikingly similar across life’s major domains: Evidence for life’s metabolic optimum. Proc Natnl Acad Sci USA 105, 16994–16999.

42. Deere JA, Chown SL. 2006 Testing the beneficial acclimation hypothesis and its alternatives for locomotor performance. Am Nat 168, 630–644.

43. Hester ET, Doyle MW. 2011 Human impacts to river temperature and their effects on biological processes: A quantitative synthesis. J Am Water Res Assoc 47, 571–587.

44. Dell AI, Pawar S, Savage VM. 2013 The thermal dependence of biological traits. Ecology 94, 1205.

45. Lurling M, Eshetu F, Faassen EJ, Kosten S, Huszar VLM. 2013 Comparison of cyanobacterial and green algal growth rates at different temperatures. Freshwater Biol 58, 552–559.

46. Ras M, Steyer J-P, Bernard O. 2013 Temperature effect on microalgae: a crucial factor for outdoor production. Rev Environ Sci Bio/Technol 12, 153–164.

47. Hinchliff CE, Smith SA, Allman JF, Burleigh JG, Chaudhary R, Coghill LM, Crandall KA, Deng J, Drew BT, Gazis R et al. 2015 Synthesis of phylogeny and taxonomy into a comprehensive tree of life. Proc Natnl Acad Sci USA 112, 12764–12769.

48. Chamberlain S. 2017 rgbif: Interface to the Global ‘Biodiversity’ Infomation Facility ‘API’. R package version 0.9.8. https://CRAN.R-project.org/package=rgbif

49. Chamberlain S, Ram K, Hart T. 2016 spocc: Interface to Species Occurrence Data Sources. R package version 0.5.0. http://CRAN.R-project.org/package=spocc

50. R Development Core Team. 2017 R: A language and environment for statistical computing. R Foundation for Statistical Computing, Vienna, Austria.

51. Hodar JA. 1996 The use of regression equations for estimation of arthropod biomass in ecological studies. Acta Oecol 17, 421–433.

52. Ganihar SR. 1997 Biomass estimates of terrestrial arthropods based on body length. J Bioscience 22, 219–224.

53. Lynch M. 1991 Methods for the analysis of comparative data in evolutionary biology. Evolution 45, 1065–1080.

54. Housworth EA, Martins EP, Lynch M. 2004 The phylogenetic mixed model. Am Nat 163, 84–96.

55. Hadfield JD, Nakagawa S. 2010 General quantitative genetic methods for comparative biology: phylogenies, taxonomies and multi-trait models for continuous and categorical characters. J Evol Biol 23, 494–508.

56. Gilmour AR, Gogel BJ, Cullis BR, Thompson R. 2009 ASReml user guide. Release 3.0 (NSW Department of Industry and Investment, Sydney, Australia).

57. R Development Core Team. 2013 R: A language and environment for statistical computing. R Foundation for Statistical Computing, Vienna, Austria.

58. Hadfield JD. 2010 MCMC methods for multi-response generalized linear models: the MCMCglmm R Package. J Statistical Softw 33, 1–22.

59. Michonneau F, Brown J, Winter D. 2016 rotl: Interface to the ‘Open Tree of Life’ API. R package version 0.5.0. https://CRAN.R-project.org/package=rotl.

60. Hahn MW, Pöckl M. 2005 Ecotypes of planktonic actinobacteria with identical 16S rRNA genes adapted to thermal niches in temperate, subtropical, and tropical freshwater habitats. Appl Environ Microbiol 71, 766–773.

61. Sorokin C, Krauss RW. 1962 Effects of temperature & illuminance on Chlorella growth uncoupled from cell division. Plant Physiol 37, 37–42.

62. Mattiuzzi M, Detsch F. 2017 MODIS: Acquisition and Processing of MODIS Products. v1.1.0.https://CRAN.R-project.org/package=MODIS.

63. Hijmans RJ. 2016 raster: Geographic Data Analysis and Modeling. v2.5-8. http://CRAN.R-64.project.org/package=raster/.

64. Ryan RA, Ulrich J. 2014 xts: eXtensible Time Series. v0.9-7. https://CRAN.R-project.org/package=xts.

65. Felsenstein J. 1985 Phylogenies and the comparative method. Am Nat 125, 1–15.

66. Grafen A. 1989 The phylogenetic regression. Phil Trans R Soc B 326, 119–157.

67. Kenward MG, Roger JH. 1997 Small sample inference for fixed effects from restricted maximum likelihood. Biometrics 53, 983–997.

68. Pagel M. 1999 Inferring the historical patterns of biological evolution. Nature 401, 877–884.

69. White I. 2013 R pin function. http://www.homepages.ed.ac.uk/iwhite//asreml/.

70. Hochachka PW, Somero GN. 2002 Biochemical Adaptation. Mechanism and Process in Physiological Evolution. Oxford University Press, Oxford.

71. Sunday JM, Bates AE, Kearney MR, Colwell RK, Dulvy NK, Longino JT, Huey RB. 2014 Thermal-safety margins and the necessity of thermoregulatory behavior across latitude and elevation. Proc Natnl Acad Sci USA 111, 5610–5615.

72. Denny MW. 1993 Air and Water: The Biology and Physics of Life’s Media. Princeton University Press, Princeton.

73. Michaletz ST, Kerkhoff AJ, Enquist BJ. 2017 Drivers of terrestrial plant production across broad geographical gradients. Glob Ecol Biogeogr 27, 166–174.

74. Hodkinson ID. 2003 Metabolic cold adaptation in arthropods: a smaller-scale perspective. Funct Ecol 17, 562–567.

75. Irlich UM, Terblanche JS, Blackburn TM, Chown SL. 2009 Insect rate-temperature relationships: environmental variation and the metabolic theory of ecology. Am Nat 174, 819–835.

76. Heinrich B. 1977 Why have some animals evolved to regulate a high body temperature? Am Nat 111, 623–640.

